# MicroED structure of the C11 cysteine protease Clostripain

**DOI:** 10.1101/2024.01.04.574240

**Authors:** Yasmeen N. Ruma, Guanhong Bu, Tamir Gonen

## Abstract

Clostripain secreted from *Clostridium histolyticum* is the founding member of the C11 family of Clan CD cysteine peptidases, which is an important group of peptidases secreted by numerous bacteria. Clostripain is an arginine specific endopeptidase. Because of its efficacy as a cysteine peptidase, it is widely used in laboratory settings. Despite its importance the structure of clostripain remains unsolved. Here we describe the first structure of an active form of *C. histolyticum* Clostripain determined at 3.6 Å resolution using microcrystal electron diffraction (MicroED). The structure was determined from a single nanocrystal after focused ion beam milling. The structure of Clostripain shows a typical Clan CD α/β/α sandwich architecture and the Cys231/His176 catalytic dyad in the active site. It has a large electronegative substrate binding pocket showing its ability to accommodate large and diverse substrates. A loop in the heavy chain formed between residues 452 to 457 is potentially important for substrate binding. In conclusion, this result demonstrates the importance of MicroED to determine the unknown structure of macromolecules such as Clostripain, which can be further used as a platform to study substrate binding and design of potential inhibitors against this class of peptidases.

## Introduction

*Clostridium histolyticum* / *Hathewaya histolytica* is a gram-positive pathogenic bacterium that is known to cause local necrosis in human muscles, organs and connective tissues. *C. histolyticum* can secrete five different kinds of potent exotoxins that includes proteinases and collagenases (Nishida and Imaizumi, 1966; Oakley and Warrack, 1950). These toxins can cause proteolysis and degradation of cells (Hatheway, 1990), thus leading to systemic toxemia (Durmaz et al., 2000) and eventually death if left untreated (Flores-Diaz and Alape-Giron, 2003). In addition to the collagenases and peptidases, a cysteine - activated protease, known as Clostripain was also isolated from the culture filtrates of *C. histolyticum* (E. Labrou, 2013; Kocholaty et al., 1938). However, there is no report on the proteolysis effect of Clostripain on human cells yet. The protease is mainly used *in vitro* as an important research tool for protein sequencing and peptide fragment condensation, e.g., its application in human islet isolation procedure (Ståhle et al., 2015).

Clostripain is the archetypal member of the C11 family of the Clan CD of cysteine endopeptidases (Rawlings et al., 2012). Since its discovery in 1938, there have been ongoing research on Clostripain (Kembhavi et al., 1991; Labrou and Rigden, 2004; Ullmann and Jakubke, 1994; Witte et al., 1996) and the homologue proteases of the C11 family (Manabe et al., 2010; McLuskey et al., 2016; McLuskey and Mottram, 2015). Although the proteolysis effect of Clostripain on pathogenic process is not yet known, there have been reports of the involvement of other C11 proteases in the pathogenicity of disease in humans. For example, a Clostripain-like protease secreted from the commensal pathogen *Clostridium perfringens* was reported to promote macrophage phagocytosis by degradation of host neutrophils (Guzik et al., 2007). Another peptidase from the same family, Fragipain from *Bacteroides fragilis* was reported to cause sepsis in mice (Choi et al., 2016), thus making this family of proteases an interesting group to explore.

The members of the Clan CD cysteine peptidases are classified mainly on similar structural features and function, rather than sequence homology (Labrou and Rigden, 2004; Rawlings et al., 2012). These proteases have a highly conserved histidine/cysteine catalytic dyad and use an active cysteine to cleave protein peptide bonds (McLuskey and Mottram, 2015). The first structure determined from this family was of PmC11, from *Parabacteroides merdae* (McLuskey et al., 2016) and was used as the model to study the structure-function relationship of C11 peptidases. Then on, crystal structures of Clostripain-like proteases from different bacteria have been determined either in unbound form (Gonzalez-Paez et al., 2019; McLuskey et al., 2016) or in complex with their inhibitors (Roncase et al., 2019; Roncase et al., 2017) by X-ray diffraction. However, Clostripain which has a high specificity for arginine and requires calcium ions for activation (Kembhavi et al., 1991; Witte et al., 1994) remained without an experimental structure probably due to its low yield in heterologous host expression systems (Manabe et al., 2010) making large scale crystal screening prohibitive.

In this study, we have determined the first experimental structure of *Clostridium histolyticum* Clostripain in its active state at 3.6 Å resolution by the cryogenic electron microscope (Cryo-EM) method, microcrystal electron diffraction (MicroED). MicroED is a robust method used to determine structures of different samples including small molecules, peptides, natural products and proteins using vanishingly small crystals (Bu and Nannenga, 2021; Mu et al., 2021; Nannenga and Gonen, 2019; Nannenga et al., 2014; Shi et al., 2016). The structure consists of a typical Clan CD α/β/α sandwich architecture and the Cys231/His176 catalytic dyad in the active site. A large electronegative cavity was identified as the substrate binding pocket. The structure allowed us to identify a loop between residues 452 and 457 that may be important for substrate binding. This study demonstrates MicroED’s ability to deliver structures that were not attainable by other methods even when sample is prohibitively limiting for large scale crystal growth screening.

## Results and Discussion

### Structure determination

Initial Clostripain crystals were formed as needles within 2 to 3 days at 20 °C in presence of 0.2 M Ammonium phosphate monobasic, 0.1 M TRIS pH 8.5, 50% v/v 2-methyl-2,4-pentanediol. These crystals were difficult to reproduce and so seeding was attempted. Seeding stock made from these crystals was used to initiate crystal growth in several different conditions. Small plate-shaped crystals were observed in 0.2 M Ammonium acetate, 0.1 M Na-citrate tribasic dihydrate pH 5.6, 30% w/v PEG 4000 within a day of seeding.

While these crystals were far too small for X-ray diffraction, they were also not suitable for MicroED analyses directly as they were too thick and the electrons could not traverse the thick crystals in the transmission electron microscope (TEM). The crystals were transferred to EM grids, plunge frozen in liquid ethane and stored in liquid nitrogen and thinned using a cryogenic focused ion beam (FIB) in scanning electron microscope (SEM) to a thickness suitable for MicroED (Martynowycz et al., 2019). With this approach the crystal lamellae ∼300nm in thickness were produced (Figure 1). The grids containing the milled lamellae were then transferred to the TEM operating under cryogenic conditions. The Thermo-Fisher EPU and Velox were used for locating the lamellae and screening diffraction quality. The MicroED data were collected in counting mode with a Thermo-Fisher Falcon 4i direct electron detector using continuous rotation method (Nannenga and Gonen, 2014) and an energy filter with slit size of 20 eV.

**Figure 1.**
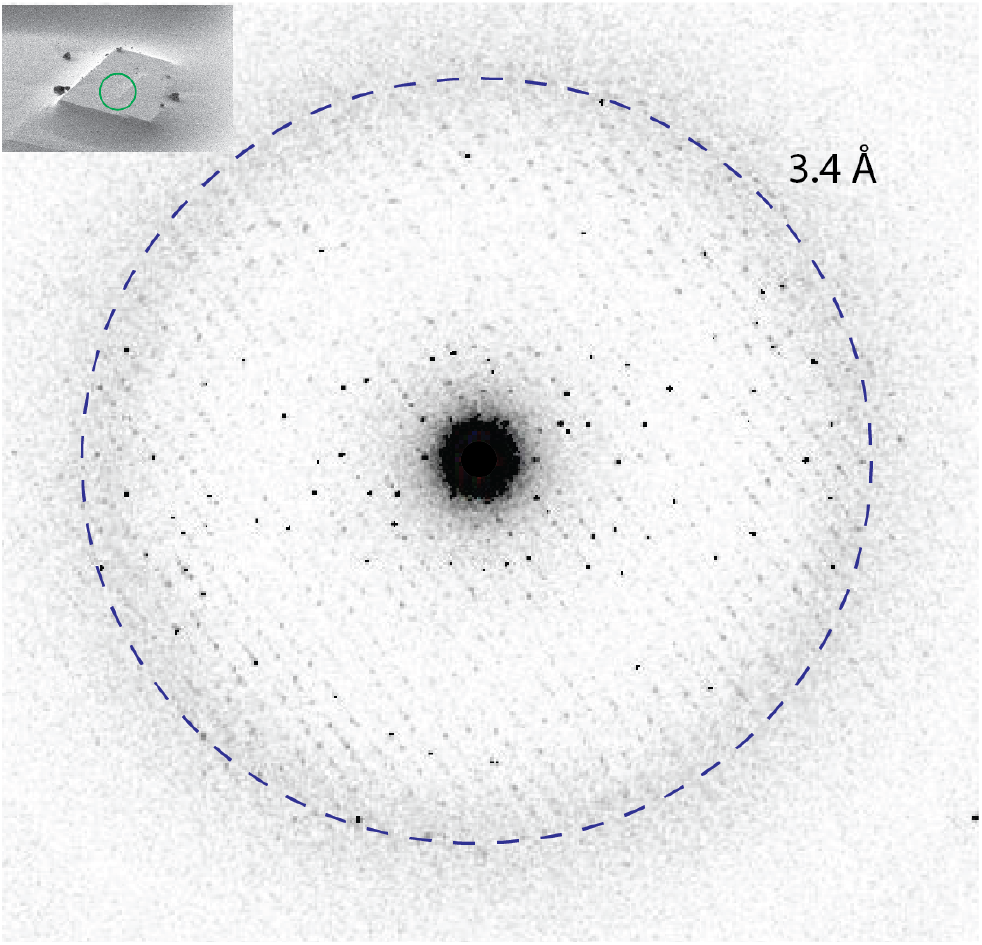
Representative MicroED diffraction pattern of a Clostripain lamella collected by continuous rotation. Inset shows a FIB/SEM image of a Clostripain crystal. Green circle represents the area used for MIcroED data collection (<1μm)

MicroED datasets were collected for a total of 6 lamellae and the dataset from the best diffracting lamella was processed (Figure 1) at a resolution of 3.60 Å and an overall completeness of 90%. The data was processed and refined as detailed in the methods section. Clostripain crystallized in the space group P22_1_2_1_ with cell dimensions of a = 63.477 Å, b = 98.92 Å, c = 145.104 Å and α = β = γ = 90°. The AlphaFold model (Model ID AF-A0A4U9RR22-F1) (Jumper et al., 2021) for *C. histolyticum* Clostripain was used to phase the data using molecular replacement. In the sequence of Clostripain, residues 1 - 27 represent the signal peptides, 28 - 50 represent the pro-peptide, 51 - 181 is the light chain, 182 - 190 is the linker joining the light and heavy chain and 191 - 526 is the heavy chain of Clostripain (Uniprot ID: P09870) (Figure 2). After the initial phasing and refinement, no densities for the signal peptide, pro-peptide and the linker were observed indicating that the MR did not result in any significant model bias from the AlphaFold model used for phasing. There was also no density seen for the residues 452 – 457 in the heavy chain suggesting that these residues form a flexible dynamic loop. For subsequent refinement these sections of the protein were removed. After several rounds of refinement and manual building/rebilding, the structure was finalized with an acceptable R_work_/R_free_ = 33%/34%. At 3.6 Å resolution, and with 91.1 % of the residues in the favorable region of the Ramachandran plot. The final model has a dimer of Clostripain heterodimer in the asymmetric unit (ASU). Each monomer comprises the light chain (residues 51-181) and the heavy chain (residues 191-526).

**Figure 2.**
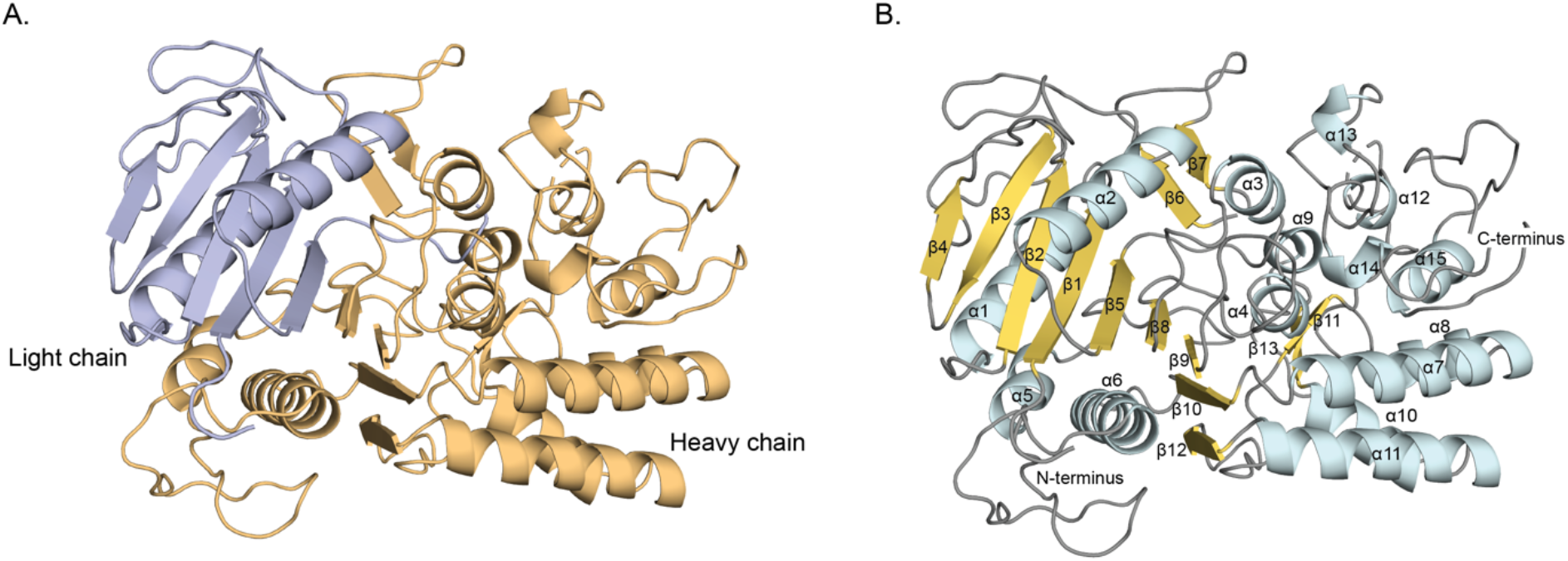
MicroED structure of Clostripain. (A) Quaternary structure of Clostripain. Light chain is shown in light purple and heavy chain is shown in light orange. (B) Tertiary structure of Clostripain. Alpha helices are shown in light cyan, beta strands in light yellow and the loops in gray. The N and C termini, alpha helices and beta strands are all labelled. The helices and strands are numbered based on the sequence starting from the N-terminus.

**Table 1.**
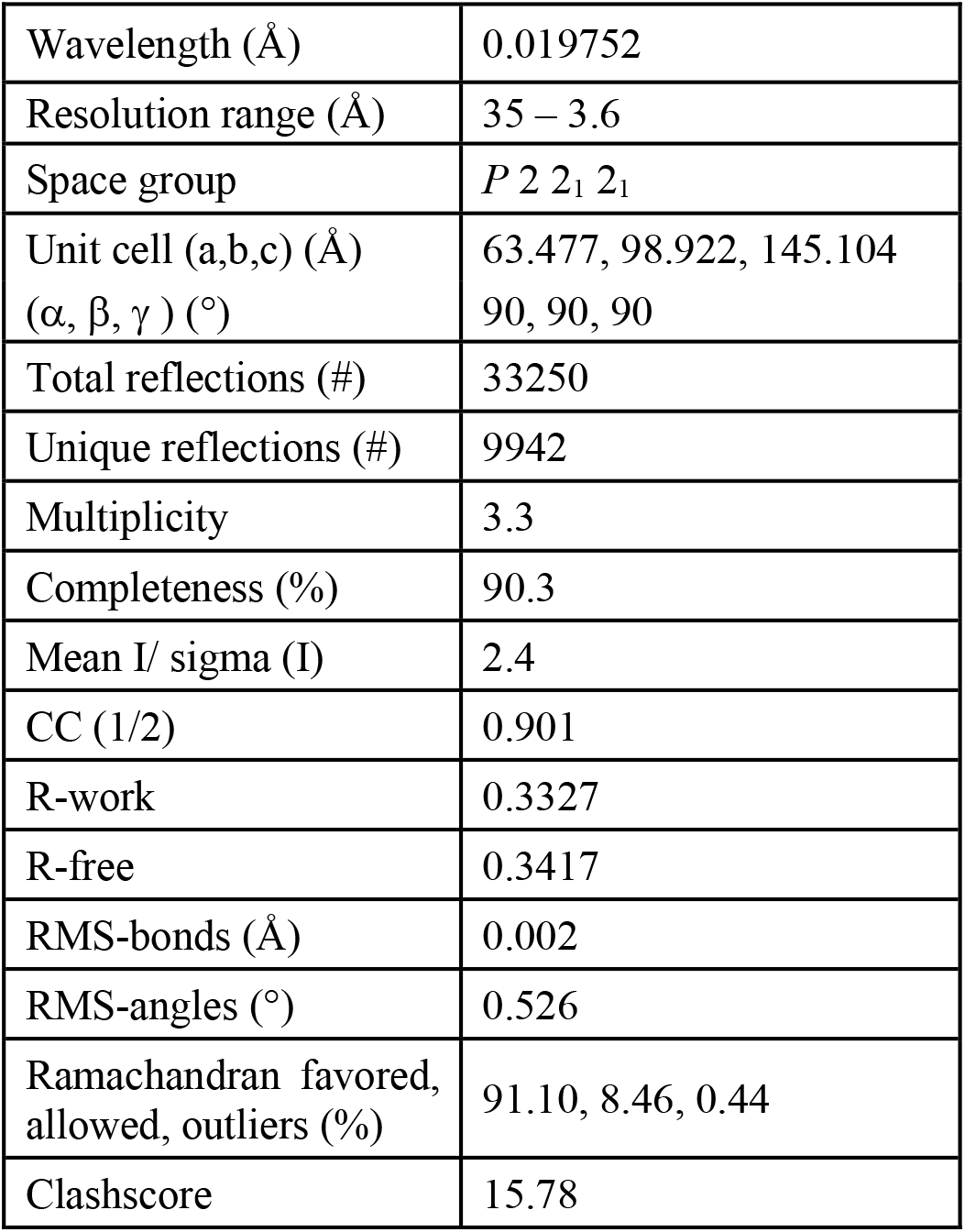
MicroED data collection and refinement statistics.

### Overall structure of Clostripain

Clostripain in its proenzyme form is a 59 kDa sized protein. Its polypeptide chain is composed of a heavy and a light chain that are connected by a linker region and held together by strong non-covalent forces. When a calcium ion activates the proenzyme, it undergoes auto-maturation by autocatalyzing the removal of the linker peptide at its two cleavage sites - Arg 181, part of the light chain (labelled in Figure 3A and 3C) and Arg 190, part of the linker peptide (Witte et al., 1996).

**Figure 3.**
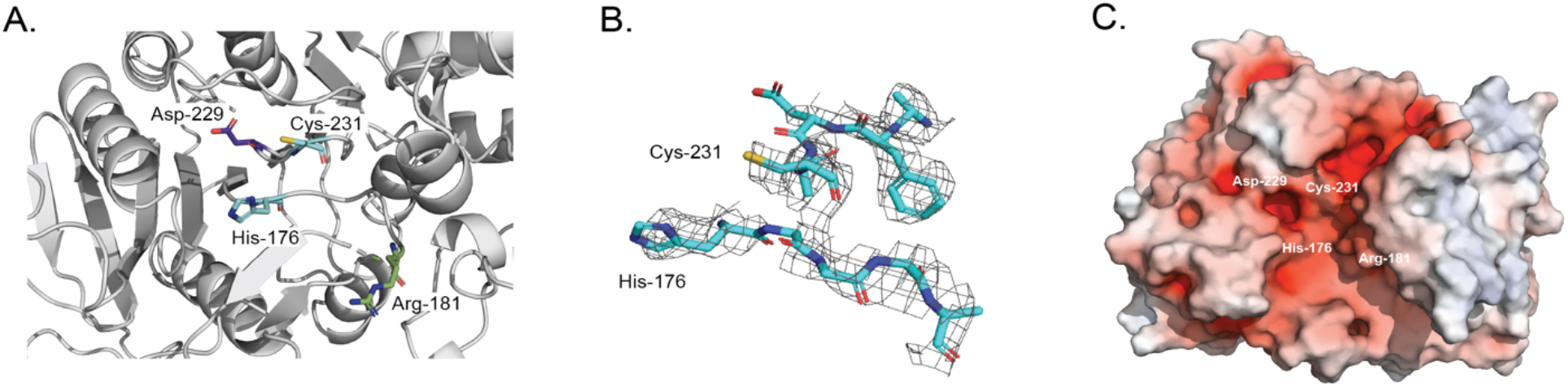
(A) Structure of Clostripain highlighting the catalytic dyad, His176 and Cys231 in cyan blue sticks. The cleavage site Arg181 is presented as green stick and the P1 specific substrate site Asp229 as purple stick. Nitrogen, oxygen and sulphur atoms are colored blue, red and yellow respectively. (B) 2Fo-Fc map (gray mesh) contoured at 1σ showing density for the active site residues. (C) Electrostatic surface potential of Clostripain in the same orientation as A showing the same residues as A, where blue and red denote positively and negatively charged surface potential, respectively, contoured at ±10 kT/e.

The residues from both the light chain and the heavy chain have been resolved in the MicroED structure (Figure 2A). The linker between the light and heavy chain could not be resolved since the protease was in its active state.

Clostripain has a typical C11 protease structure as reported for other Clostripain-like proteases (Gonzalez-Paez et al., 2019; McLuskey et al., 2016; Roncase et al., 2019; Roncase et al., 2017). Overall, Clostripain consists of 15 alpha helices and 13 beta strands, with 2 alpha helices and 5 beta strands in the light chain and 13 helices and 8 beta strands in the heavy chain (Figure 2B). The alpha helices and beta strands are numbered starting from the N-terminal region at residue 51 and ending in 526 at the C-terminal (Figure 2B). The structure is made up of a central nine-stranded β-sheet typical of C11 proteases (Gonzalez-Paez et al., 2019; McLuskey et al., 2016; Roncase et al., 2019; Roncase et al., 2017) forming an α/β/α sandwich architecture (Figure 2B). The β-strands involved are β1 - β5, β8 - β10 and β12. Out of the 9 beta strands, 7 β-strands are parallel and 2 strands are antiparallel (β3 and β10) (Figure 2B). Among the 9 β-strands, β1 - β5 belong to the light chain, and the rest three are of the heavy chain (Figure 2). The alpha helices surrounding the beta sheet includes α1 (from light chain), α5 and α6 (from heavy chain) on one side and α2 (light chain) and α4 (heavy chain) on the other side. There are two pairs of β-hairpins (β6 and β7, and β11 and β13) antiparallel to each other.

The active site of Clostripain (Figure 3A) consists of a catalytic dyad with the residues His176 and Cys231 (Labrou and Rigden, 2004). His176 belongs to the light chain and Cys231 belongs to the heavy chain, with clear density for these residues resolved in the map (Figure 3B).

Electrostatic surface analysis (Figure 3C) shows that the catalytic dyad forms part of a large electronegative pocket consistent with a binding site for a positively charged substrate, confirming its arginine specific cleavage and its potential to accommodate larger peptides. The pocket is also lined with the residue Asp229 (shown in Figure 3A and 3C) which has been predicted to be the P1 specificity determining residue in Clostripain (Ullmann and Jakubke, 1994), similar to Asp177 shown in PmC11 previously (McLuskey et al., 2016).

### Comparison of the active Clostripain structure with the AlphaFold model representing the inactive form

Superposition of the experimental MicroED structure with the predicted AlphaFold model highlights important functional differences. Superposition of the two structures (Figure 4A) revealed that the structures differed overall with an r.m.s.d of 0.533 Å. Strikingly, the AlphaFold model predicts the proenzyme structure while we determined the activated enzyme structure (Figure 2). Comparison of the proenzyme model with the experimental structure of the active protease highlights the effect of the linker peptide on the substrate binding pocket. The electrostatic surface diagram of the proenzyme demonstrated that the linker between the light and heavy chain has an electropositive surface, and it gates the substrate binding pocket (Figure 4B). Once the enzyme is activated, the linker is cleaved out therefore exposing the substrate binding pocket (Figure 4C) to peptides. The removal of the gating peptide also changes the overall electrostatics making them more favorable for substrate binding.

**Figure 4.**
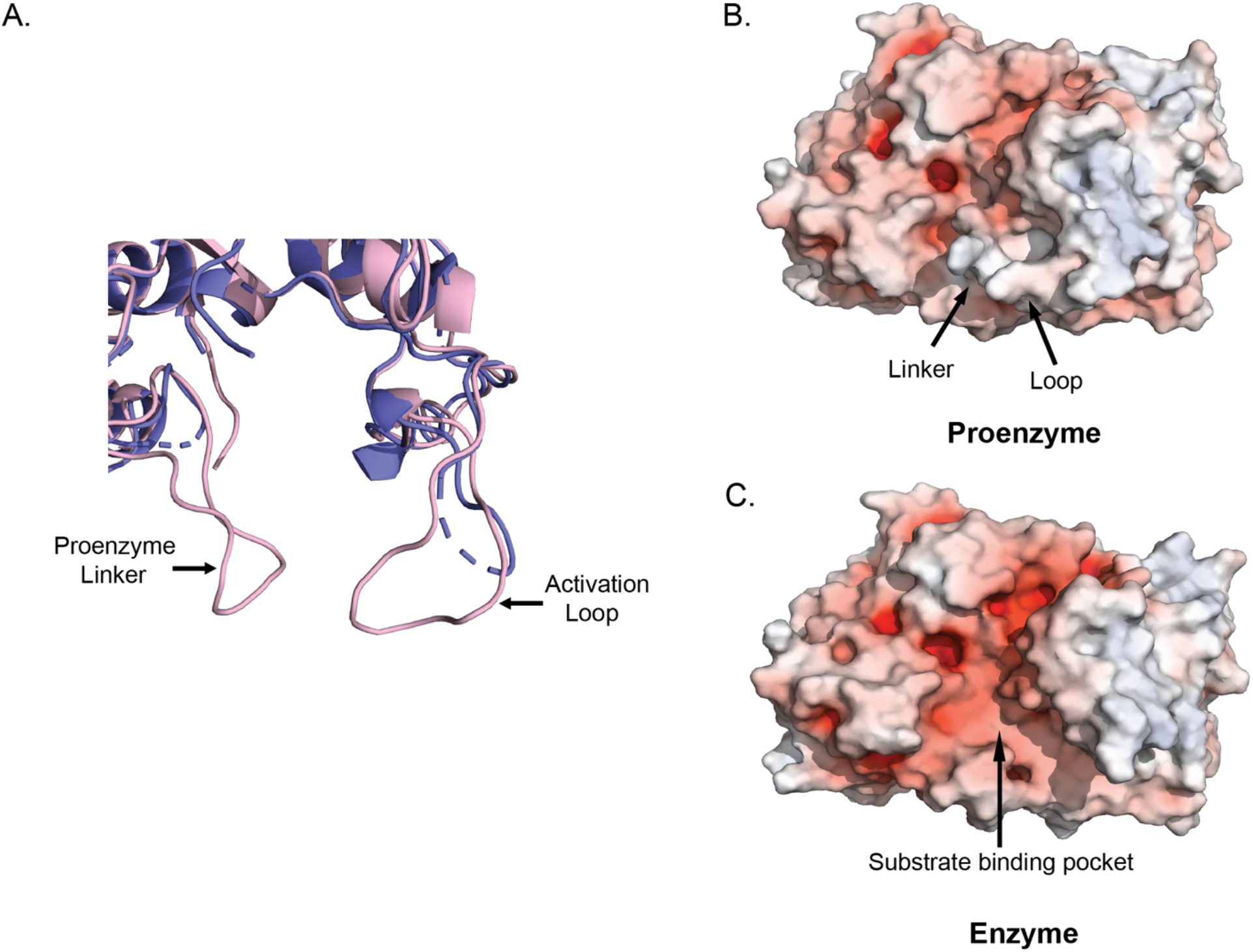
(A) Superposition of the MicroED structure (purple cartoon) and the AlphaFold model (light pink cartoon) of Clostripain highlighting the linker and the loop. The linker and the loop formed by residues 452 – 457 are labelled on the proenzyme. (B) Electrostatic surface for the AlphaFold model representing the proenzyme. (C) Electrostatic surface for the active Clostripain MicroED structure. Here blue and red denote positively and negatively charged surface potential, respectively, contoured at ±10 kT/e.

Furthermore, we hypothesize that the loop formed by the residues 452 to 457 (STYYTS), (circled in Figure 4A) plays a significant role in substrate binding. In the proenzyme, it is noticed that the loop projects towards the active site and provides support to the linker to keep the active site closed (Figure 4B). So, absence of density for these residues in the MicroED structure could potentially mean that when the protease is activated and linker removed, this loop becomes more dynamic and undergoes a structural rearrangement to accommodate the binding of substrates. A similar structural mechanism was previously described in the structure of metacaspase from *Trypanosoma brucei* (PDB ID - 4AF8), a C14 clan CD peptidase, which also requires Calcium ion for activation and is arginine specific. In presence of Calcium ion, it was shown that a conformational change occurs in loop 7 of the protease thus facilitating substrate binding (McLuskey et al., 2012). Consistent with the above reports this suggests to us that a similar mechanism may exist in Clostripain.

### Structural comparison of Clostripain with other C11 Proteases structures in PDB

The structure of Clostripain was compared with other C-11 protease’s structure by a DALI structural similarity search (Holm and Rosenstrom, 2010). The top related structures include: Clostripain-related protein from *B. thetaiotaomicron* (PDB ID - 6N9J, Z score = 33.3, rmsd = 2.4, number of residues = 314/356, % ID = 20) (Roncase et al., 2019), inactive zymogen C11 protease from *Parabacteroides distasonis* (PDB ID - 6MZO, Z score = 32.2, rmsd = 2.7, number of residues = 311/346, %ID = 21 (Gonzalez-Paez et al., 2019), Cysteine protease from *B. fragilis* (PDB ID - 5DYN, Z score = 18.5, rmsd = 2.6, number of residues = 220/245, %ID = 19) (Choi et al., 2016), Protease from *Parabacteroides merdae* (PDB ID – 3UWS, Z score =18.2, rmsd = 2.6, number of residues = 216/228, % ID = 20)(McLuskey et al., 2016), Cysteine protease Gingipain from *Porphyromonas gingivalis* (PDB ID – 1CVR, Z score = 11.4, rmsd = 3.7, number of residues = 179/433, %ID =9) (Eichinger et al., 1999).

The structure of Clostripain was superposed with the two structures of highest similarity (PDB ID 6N9J and 6MZ0) as shown in Figure 5. Overall, the structures share similar central α/β/α sandwich architecture. As seen in Figure 5A, the active site residues His176 and Cys231 of Clostripain superimpose with His134 and Cys183 and have highly conserved spatial orientation. The distance between Cys-His of Clostripain is 5.1 Å close to that of 6 Å in the protease from *B. thetaiotaomicron*. Overlapping with the inactive zymogen protease (Figure 5B) shows similar structural features, except that the His residue, His135 in 6MZO is closer to the core (Gonzalez-Paez et al., 2019) and orients differently to His176 in Clostripain. The distance between the residues is 5.7 Å, comparable to that of 5.1 Å in Clostripain.

**Figure 5.**
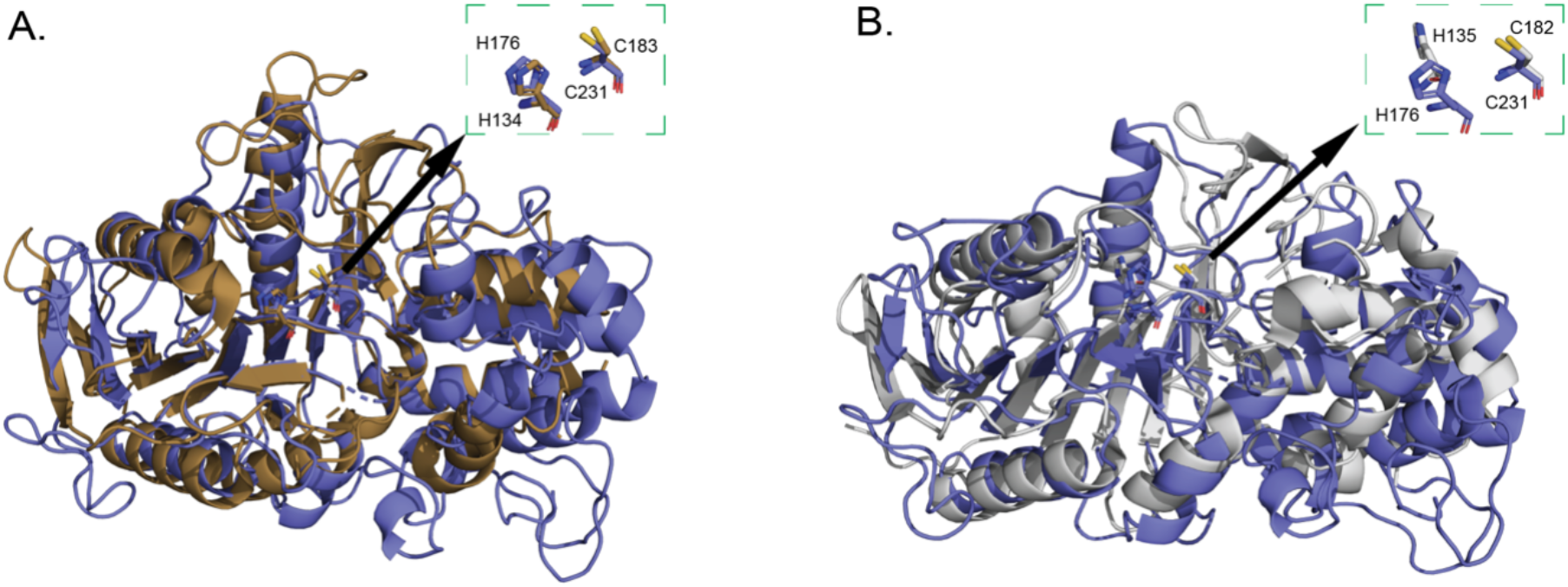
Superposition of Clostripain (purple cartoon) with (A) Clostripain-related protein from *B. thetaiotaomicron*, PDB ID – 6N9J (Brown cartoon), (B) inactive zymogen C11 protease from *Parabacteroides distasonis*, PDB ID – 6MZO (Gray cartoon). The active site residues are shown in the insets.

## Conclusion

Here we report the previously unknown structure of a C11 protease, Clostripain from *C. histolyticum* in its active form determined by MicroED. While the structure of Clostripain was unattainable by other methods, MicroED delivered a 3.6 Å resolution structure using a single nanocrystal. This study adds to the growing list of novel macromolecular structures determined by MicroED and can form the basis for future development of new protease inhibitors. Future work on substrate binding in a time resolved manner may provide additional insight into the mechanistic basis of substrate specificity and mode of activation of this protease.

## Materials and methods

### Materials

Clostripain (from *C. histolyticum*) in lyophilized, pre-activated form was purchased from Abnova (Taiwan), and used without further purification. Crystallization screens were purchased from Hampton Research (Aliso Viejo, CA). All reagents were made with MilliQ water. TBS was purchased from BioRad.

### Crystallization

Clostripain was solubilized in 1x TBS at a concentration of 10 mg/ml for setting up crystallization plates. Crystallization screens were set by sitting drop vapor diffusion method using a Mosquito crystallization robot, in 200 nl drops, at 1:1 sample to mother liquor ratio in 96-well intelliplates (Hampton research). Initial needle-shaped crystals were formed within 2 to 3 days at 20 °C in presence of 0.2 M Ammonium phosphate monobasic, 0.1 M TRIS pH 8.5, 50% v/v 2-methyl-2,4-pentanediol. Seeding stock was prepared using the protocol specified in the seed bead steel kit HR4-780 (Hampton research). The seeding stock was used to make robust crystals in 24 well plates by hanging drop vapor diffusion in the mixture of 1 μl protein, 0.1 μl the seeding stock and 1 μl crystallization condition of 0.2 M Ammonium acetate, 0.1 M Na-citrate tribasic dihydrate pH 5.6, 30% w/v PEG 4000. The plates were kept at 20 °C and crystal growth was observed in 1 day.

### Sample preparation

The EM grids with Clostripain crystals were prepared in a Leica GP2 plunge freezer set to 95% humidity and 20 °C temperature as described previously (Martynowycz et al., 2019). Quantifoil Cu 200 R2/2 holey carbon grids (Quantifoil) were glow discharged negatively for 45 sec before sample application. The crystal drops were then diluted with 2 μl of crystallization condition and then applied to the grids. The grids were each blotted for 20 sec plunge-frozen into liquid ethane. The grids were stored in liquid nitrogen until use. The grids were clipped prior to screening in the electron microscope.

### FIB milling of the crystals

Clipped grids containing Clostripain crystals were then loaded into a Aquilos dual-beam FIB/SEM (Thermo Fisher) operating at -180 °C following procedures described previously (Martynowycz et al., 2019). The grids were sputter coated with a thin layer of platinum to preserve the sample during imaging and ion beam milling. Complete atlases of the grids were acquired using the MAPS software (Thermo Fisher). The milling sites were located and the eucentric height adjusted. The crystals were then milled using a gallium ion beam. The current used for milling was gradually reduced from 0.5 nA to 30 pA with every reduction in the thickness of the lamella. Milling was stopped when the lamella of desired thickness 300 nm was achieved.

### MicroED Data Collection

After milling, the grids with the milled lamella were transferred to Titan Krios G3i TEM (Thermo Fisher) operating at -190 °C an accelerating voltage of 300 kV (∼0.019687 Å wavelength). The Krios is equipped with a field emission gun and a Falcon 4i direct electron detector. The software EPU (Thermo Fisher) was used to acquire a low magnification atlas of the whole grid to identify the lamella. The stage position was moved to the lamella and the eucentric height then adjusted by taking live view in Velox (Thermo Fisher). A selectris energy filter operating at a slit width of 20 eV was used for the data collection. The selected area (SA) aperture (∼2 μm in diameter) was inserted and centered on the desired area to obstruct any background reflections. Initial screening of the lamella for diffraction was carried out using Velox. Lamella showing quality diffraction spots were used for data collection. MicroED data were collected by continuous rotation at a rate of 0.3 °/s for 420 sec, with a wedge of 90° using SerialEM. The sample to detector distance was set to the calibrated distance of 2,941 mm. The data were collected using continuous rotation method with a Falcon 4i direct electron detector in counting mode in EER format (Martynowycz et al., 2022).

### MicroED Data Processing

The diffraction data obtained in EER format was then converted to SMV format using MicroED tools (https://cryoem.ucla.edu/downloads/snapshots) (Hattne et al., 2015; Martynowycz et al., 2019). The diffraction dataset was indexed and integrated in XDS (Kabsch, 2010). Integrated intensities were scaled using in XSCALE (Kabsch, 2010). Molecular replacement was carried out in PHASER (McCoy et al., 2007) using the AlphaFold model (A0A4U9RR22) available in the AlphaFold database as the template. Structure refinement and modelling were carried out in phenix.refine (Afonine et al., 2012) and Coot (Emsley and Cowtan, 2004) respectively. Pymol (Schrodinger) was used to generate figures. Figures were assembled in Adobe illustrator.

## Data availability

Coordinates and maps were deposited in the protein data bank (Accession code XXXX) and the EM Data bank (Accession code YYYY).

## Acknowledgments

This study was supported by the National Institutes of Health P41GM136508. The Gonen laboratory is supported by funds from the Howard Hughes Medical Institute.

